# Protein farnesylation is upregulated in Alzheimer’s human brains and neuron-specific suppression of farnesyltransferase mitigates pathogenic processes in Alzheimer’s model mice

**DOI:** 10.1101/2021.05.19.444846

**Authors:** Angela Jeong, Shaowu Cheng, Rui Zhong, David A. Bennett, Martin O. Bergö, Ling Li

## Abstract

The pathogenic mechanisms underlying the development of Alzheimer’s disease (AD) remain elusive and to date there are no effective prevention or treatment for AD. Farnesyltransferase (FT) catalyzes a key posttranslational modification process called farnesylation, in which the isoprenoid farnesyl pyrophosphate is attached to target proteins, facilitating their membrane localization and their interactions with downstream effectors. Farnesylated proteins, including the Ras superfamily of small GTPases, are involved in regulating diverse physiological and pathological processes. Emerging evidence suggests that isoprenoids and farnesylated proteins may play an important role in the pathogenesis of AD. However, the dynamics of FT and protein farnesylation in human brains and the specific role of neuronal FT in the pathogenic progression of AD are not known. Here, using postmortem brain tissue from individuals with no cognitive impairment (NCI), mild cognitive impairment (MCI), or Alzheimer’s dementia, we found that the levels of FT and membrane-associated H-Ras, an exclusively farnesylated protein, and its downstream effector ERK were markedly increased in AD and MCI compared with NCI. To elucidate the specific role of neuronal FT in AD pathogenesis, we generated the transgenic AD model APP/PS1 mice with forebrain neuron-specific FT knockout, followed by a battery of behavioral assessments, biochemical assays, and unbiased transcriptomic analysis. Our results showed that the neuronal FT deletion mitigates memory impairment and amyloid neuropathology in APP/PS1 mice through suppressing amyloid generation and reversing the pathogenic hyperactivation of mTORC1 signaling. These findings suggest that aberrant upregulation of protein farnesylation is an early driving force in the pathogenic cascade of AD and that targeting FT or its downstream signaling pathways presents a viable therapeutic strategy against AD.

## Introduction

Alzheimer’s disease (AD) is the most common cause of age-related dementia, characterized by progressive cognitive decline, and widespread deposition of neuritic plaques and neurofibrillary tangles in the brain. However, the pathogenic mechanisms underlying the development of AD remain elusive. In recent years, genome-wide association studies have identified several other genes besides APOE that are involved in cholesterol metabolism as top genetic risk factors for late-onset AD, generating much attention to the importance of cholesterol-related pathways in AD pathogenesis [1-3]. Cholesterol is one of many products of the mevalonate pathway (**Figure 1**), and farnesyl pyrophosphate (FPP) serves as a branch point for cholesterol and nonsterol isoprenoids synthesis [4]. Emerging evidence suggests that these isoprenoids play important roles in the development of AD [5-8]. Isoprenoids, including FPP and geranylgeranyl pyrophosphate (GGPP), are short-chain lipid molecules. Three enzymes, farnesyltransferase (FT), geranylgeranyl transferase-1 (GGT-1; GGT), and - 2 (GGT-2, also known as RabGGT) covalently attach FPP or GGPP to a cysteine residue in the C-terminal CAAX motif of target proteins [9] (**Figure 1**). These posttranslational modification processes, namely farnesylation and geranylgeranylation, are collectively called protein prenylation [10]. Attachment of isoprenoids enhances the lipophilicity at the C-terminus, facilitates the anchoring of the proteins to endomembrane/plasma membranes, and subsequently enables their interactions with downstream effectors. The superfamily of small GTPases, including Ras, Rho, and Rab GTPases, is the most widely studied group of proteins that are modified by prenylation. These small GTPases and their downstream signaling pathways regulate a wide range of cellular processes such as regulation of cell cycle entry, endosomal trafficking, cell adhesion and migration [11-13]. Dysregulation of protein prenylation and dysfunction of small GTPases have been implicated in various disorders, including progeria and neurodegenerative diseases, as well as cancers and cardiovascular or cerebrovascular diseases [7, 8, 14].

**Figure 1.**
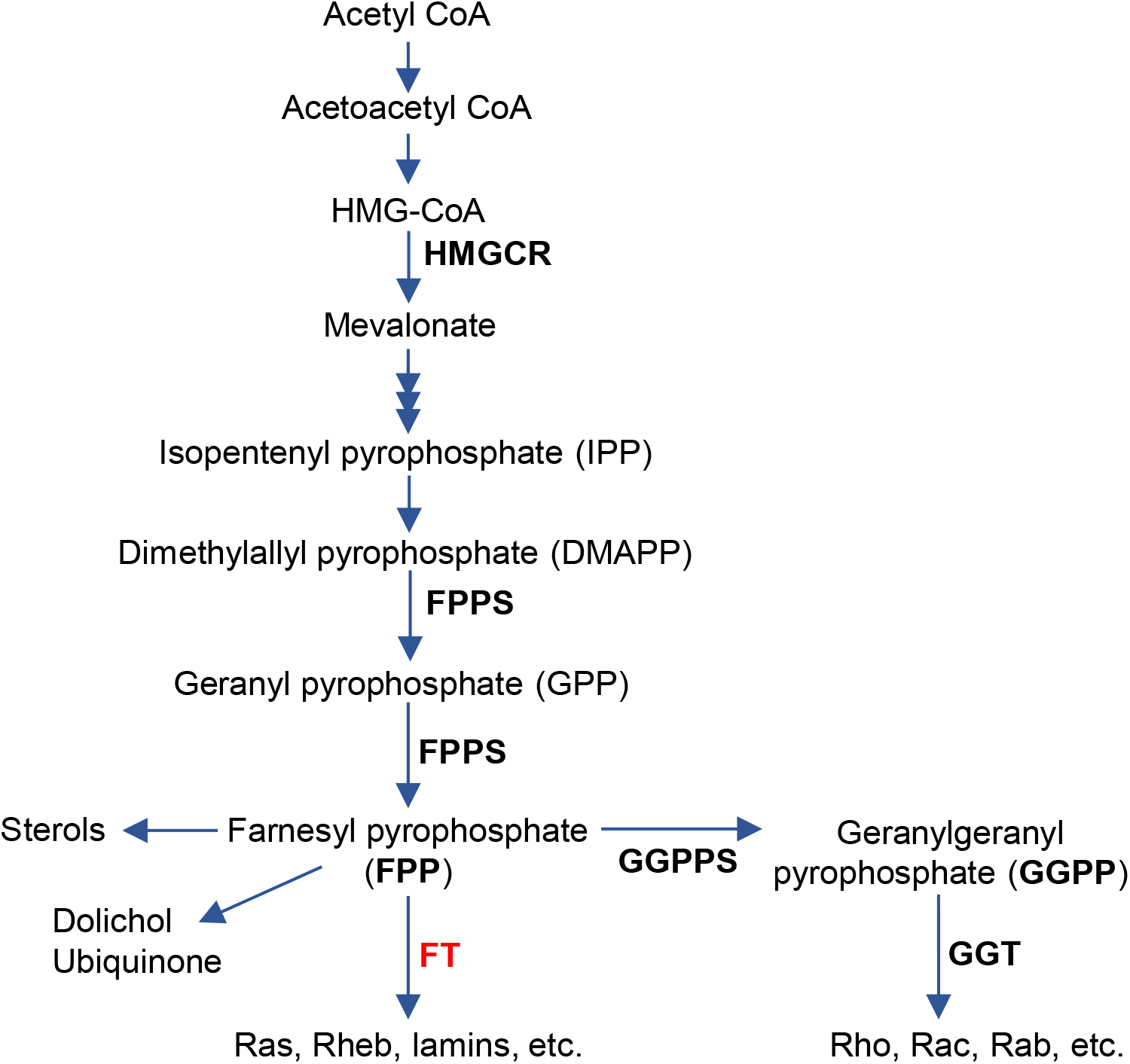
The mevalonate/isoprenoid and protein prenylation pathways. Farnesyl pyrophosphate (FPP) is derived from mevalonate, serving as the branching point for the non-steroid isoprenoid pathway and the sterol synthesis pathway. FPP and geranylgeranyl pyrophosphate (GGPP) are used for protein farnesylation and geranylgeranylation (collectively called protein prenylation) by the catalytic actions of farnesyltransferase (FT) and geranylgeranyl transferases (GGTs), respectively. HMG-CoA: 3-Hydroxy-3-methylglutaryl-CoA, HMGCR: Hydroxymethylglutaryl-CoA reductase, GPPS: Geranyl pyrophosphate synthase, FPPS: Farnesyl pyrophosphate synthase, GGPPS: Geranylgeranyl pyrophosphate synthase, FT: Farnesyltransferase, GGT: Geranylgeranyltransferase-1/2.

Pharmacological inhibition of isoprenoid synthesis using statins [15-18], protein farnesylation using FT inhibitors [19, 20], or inhibition of downstream signaling cascades of Ras and Rho GTPases [19, 21, 22] has shown benefits against amyloid-β (Aβ)/tau pathologies and synaptic/cognitive dysfunction in cellular and transgenic animal models. In humans, the levels of FPP and GGPP are elevated in the brain of individuals with AD [23, 24], and the overactivation of Ras/Rho GTPases signaling cascade correlates with the progression of amyloid or tangle pathologies [25, 26]. Collectively, these lines of evidence suggest that dysregulation of isoprenoid synthesis and overactivation of Ras/Rho GTPases pathways influence the pathogenesis of AD. However, the dynamics of protein prenylation *per se* in human brains is unknown and the specific roles of neuronal prenyltransferases are not understood. Further, pharmacological manipulations may be compromised by off-target effects, and genetic approaches that directly target the prenyltransferases are required to unravel the role of protein prenylation in the pathogenic process of AD.

Previously, we have shown that systemic haplodeficiency of either FT or GGT reduces Aβ deposition and neuroinflammation; however, intriguingly, only FT haplodeficiency rescues cognitive deficits in the APP/PS1 transgenic mouse model of AD [27], suggesting distinct roles of FT and GGT in regulating neuronal function. In follow-up studies, we found that GGT haplodeficiency significantly impairs synaptic and cognitive function whereas FT haplodeficiency has no adverse effects under physiological conditions [28, 29], indicating FT as a safer therapeutic target than GGT. Here, using postmortem brain tissue samples from individuals at different stages of AD, namely no cognitive impairment (NCI), mild cognitive impairment (NCI), or Alzheimer’s dementia, we show that FT, but not GGT, is upregulated in AD brains. Correspondingly, an exclusively farnesylated protein, H-Ras, is elevated in its membrane-associated form, and its downstream effector ERK is over-activated in the brain of individuals with MCI as well as AD. To elucidate the direct impact of neuronal FT on the pathogenic process of AD, we generated forebrain neuron-specific FT knockout in APP/PS1 mice, followed by a battery of behavioral assessments, biochemical assays, and unbiased transcriptomic analysis. Our results demonstrate that the forebrain neuron-specific FT deletion improves memory function and reduces brain Aβ load in APP/PS1 mice through suppressing APP processing and reversing the pathogenic hyperactivation of mTORC1 signaling. These findings suggest that aberrant upregulation of FT and subsequent protein farnesylation is an early driving force in the pathogenic cascade of AD and that overactivation of downstream signaling pathways contributes to the development of cognitive impairment and neuropathology. Therefore, targeting FT and/or its downstream effectors presents a promising approach to mitigate neuropathology and preserve memory function in AD.

## Materials and Methods

### Human brain tissue samples

Frozen human dorsolateral prefrontal cortex (Brodmann area 9/46) samples from participants of Religious Orders Study (ROS) were obtained from Rush Alzheimer’s Disease Center, Chicago, IL. Details on the overall study design as well as the antemortem and postmortem neuropathological assessment procedures of ROS were reported previously [30]. The study was approved by an Institutional Review Board of Rush University Medical Center, and all participants signed informed consent, an Anatomic Gift Act, and a repository consent to allow their tissue and data to be shared. Brain samples from 60 participants were obtained, with 20 samples from each of three different cognitive diagnosis groups—no cognitive impairment, mild cognitive impairment, and Alzheimer’s dementia, independent of neuropathological indices, as previously described [31, 32]. Mean age at death, education level, post-mortem interval (PMI), and sex were matched among the three groups as closely as possible upon sample selection (**Table 1**).

**Table 1.** Human participant demographics.

|  | **NCI**<br>(No cognitive<br>impairment) | **MCI**<br>(Mild cognitive<br>impairment) | **AD**<br>(Alzheimer's<br>dementia) |
| --- | --- | --- | --- |
| # Samples | 20 | 20 | 20 |
| Age at Death (yr) | 87.16 ± 3.39 | 87.42 ± 3.32 | 87.38 ± 3.58 |
| Male/Female | 4/16 | 4/16 | 4/16 |
| Education (yr) | 17.60 ± 1.39 | 17.60 ± 1.39 | 17.70 ± 1.49 |
| Post-mortem Interval (hr) | 6.21 ± 3.96 | 9.31 ± 3.96 | 6.97 ± 3.96 |
| Total amyloid-β (% area) | 4.27 ± 3.76 | 1.90 ± 1.80 | 3.92 ± 3.27 |
| Tau tangle density | 3.88 ± 0.55 | 3.61 ± 0.98 | 10.47 ± 10.74 |

### Animals

APP/PS1 double transgenic mice (B6C3-Tg (APPswe, PSEN1ΔE9) 85Dbo/J; stock# 004462) were purchased from The Jackson Laboratory (Bar Harbor, ME). To generate forebrain neuron-specific FT knockout, mice homozygous for floxed alleles for the FT β-subunit (FT^f/f^) [33] were bred with the CamKIIα-Cre recombinase mice [34] to produce FT^f/+^ Cre+ mice, followed by further crossing with FT^f/f^ to generate homozygous forebrain neuron-specific FT-knockout mice (FT^f/f^Cre+). Meanwhile, APP/PS1 mice were bred with FT^f/f^ mice in a similar manner to generate APP/PS1 mice harboring homozygous floxed FT (APP/PS1/FT^f/f^). Breeding APP/PS1/FT^f/f^ mice with FT^f/f^Cre+ generated littermates with four different genotypes: APP/PS1/FT^f/f^Cre+ (APP/PS1/FTnKO), APP/PS1/FT ^f/f^Cre-(APP/PS1), FT^f/f^ Cre+ (FTnKO), and FT^f/f^Cre-(WT). Weanlings were PCR genotyped with DNA extracted from tail/ear biopsies. Both males and females were included. Behavioral assessments were performed at 9 months of age and brain samples were collected at 10 months of age for biochemical analyses. All animal procedures used for this study were prospectively reviewed and approved by the Institutional Animal Care and Use Committee of the University of Minnesota (IACUC protocol # 1908-37310A).

### Assessment of mouse behavioral function

Three behavioral tests were performed in a consecutive order to assess AD-related behaviors: the open field test (days 1-3) for exploratory behavior in response to environmental stimuli, the elevated plus-maze test (days 4 and 5) for anxiety, and the Morris water maze test (days 6–11) for spatial learning and memory. All procedures have been previously described using the ANYMAZE system and instruments (San Diego Instruments, San Diego, CA) [27, 35, 36]. Briefly, during the open field test, mice were placed in an open square box and allowed to explore for 5 minutes. The path length and the time spent in the central zone were recorded. For the elevated plus-maze, mice were allowed to explore the two open and closed arms for 5 minutes while the path length and the time spent in the open arms were recorded. The Morris water maze test was performed using a round basin filled with water surrounded by external visual cues for orientation. The testing paradigm included the acquisition learning phase with 4 trials daily for 5 consecutive days (hidden platform), followed by the probe trial for memory retention the day after (no platform), and then the visible platform trial to assess visual acuity. The escape latency and swim path length were recorded during all trials.

### Preparation of brain tissue homogenates, and membrane and cytosolic fractions

Post-mortem human brain tissue samples (∼ 50 mg) were homogenized in 300 μl low osmotic lysis buffer (5mM Tris-HCl pH 7.4, 2 mM EDTA) supplemented with protease inhibitors and phosphatase inhibitors. An aliquot was homogenized further in the equal volume of 2x Laemmli SDS buffer, sonicated, and centrifuged at 13,000 x g for 15 minutes. The supernatant was saved as the total brain tissue homogenate for immunoblot analysis. Another aliquot of the homogenate was centrifuged at 1,500 x g for 15 minutes at 4°C and the supernatant was then centrifuged at 100,000 x g for 1 h at 4°C in a Beckman Coulter Optima™ MAX-UP Ultracentrifuge using the TLA-120.1 fixed-angle rotor. The resulting supernatant was collected as the cytosolic fraction. The pellet was washed and re-suspended in the equal initial volume of lysis buffer as the membrane fraction. Both cytosol and membrane fractions were added with 5x Laemmli SDS sample buffer and equal volumes of the fractions were subjected to immunoblot analysis.

Mouse brain tissue samples were collected and processed as described previously [27, 37, 38]. Briefly, mice were deeply anesthetized and transcardially perfused with ice-cold PBS. Brains were harvested and cut sagittally into left and right hemispheres. The left hemisphere was fixed in 4% paraformaldehyde for immunohistochemical analysis. The cortex, hippocampus, and cerebellum were dissected from the right hemisphere, snap-frozen in liquid nitrogen, and stored at −80°C until processed for immunoblot analysis and ELISA.

### Cell culture and treatment

Human neuroblastoma SH-SY5Y cells stably transfected with the wild-type APP695 construct or empty (mock) pCEP4 mammalian expression vector (a kind gift from Dr. Gunter Eckert, University of Giessen, Germany) were cultured as previously described [39]. To study the effect of a farnesyltransferase inhibitor, cells were incubated in the growth media added with 1 μM tipifarnib (Sellect cat# S1453) or vehicle (dimethyl sulfoxide, DMSO) for 48 hours. Cells were harvested in PBS containing 1% triton x-100 supplemented with protease inhibitor and phosphatase inhibitor cocktails, lysed by sonication, and subjected to immunoblot analysis.

### Immunoblot analysis

The procedures for immunoblot analyses have been described previously [27, 37, 38]. Briefly, total brain tissue and cellular lysates or the cytosolic and membrane fractions were subjected to 12% or 15% SDS-PAGE under the reducing condition, and proteins were transferred to PVDF membranes. Primary and secondary antibodies used are listed in **Table 3**. The membranes were treated with the Western Lightning Plus ECL detection reagent (PerkinElmer Life Sciences, catalog no. NEL103001EA) or Clarity Western ECL substrate (Bio-Rad). Signals were captured by exposing to HyBlot CL films (Denville, catalog no. E3018) or using the iBright™ imaging system (Invitrogen). ImageJ software was used for densitometry analysis.

### Amyloid-β ELISA

Frozen mouse cortical and hippocampal samples were processed as described previously [27, 37, 38]. Briefly, brain samples were homogenized sequentially in carbonate and guanidine buffers to obtain the carbonate-soluble and carbonate-insoluble (guanidine-soluble) fractions. The levels of Aβ in these fractions were determined with Aβ40- and Aβ42-specific ELISA kits (KHB3481 and KHB3441, ThermoFisher) following the manufacturer’s protocol.

### Immunohistochemical analysis

Protocols for immunohistochemical analysis have been described previously [27, 37, 38]. Briefly, mouse brain hemispheres prefixed with 4% PFA for 48 hours were sectioned at 50 μm using a Vibratome (Leica Microsystems Inc). Immunostaining of free-floating sections was performed using the ABC kit (Vector Laboratories, Burlingame, CA). The primary antibody 6E10 was used for assessing Aβ deposition, and IBA-1 antibody was used for assessing activated microglia. Detailed information on antibodies is available in **Table 3**. Immunoreactivity of Aβ and IBA-1 in cortical and hippocampal areas was quantified using Image-Pro Plus (MediaCybernetics, Rockville, MD) and Fiji/Image J.

### RNA extraction, library preparation, and sequencing

Eighteen mice (9-10 months) were used for transcriptome analysis of the brain: 6 APP/PS1/FTnKO, 6 APP/PS1, and 6 WT controls. The Aurum™ Total RNA Fatty and Fibrous Tissue Kit (732-6830; Bio-Rad) was used to extract total RNA from the frontal cortex following the manufacturer’s instructions. RNA quantity was measured using Quant-iT RiboGreen Assay Kit (ThermoFisher), and the quality was determined by the Agilent Bioanalyzer system. An RNA integrity number (RIN) value of 8.0 or greater (marginal 7.6-7.9) was used for the RNA library preparation. RNA-seq libraries were prepared using Illumina TruSeq Stranded mRNA Kit and sequenced using Illumina NovaSeq 6000 platform, generating 20 million reads per sample on a 150-bp paired-end run. All quality control, library preparation and sequencing were conducted at the University of Minnesota Genomics Center.

### RNA sequencing read mapping and data analysis

The raw reads were trimmed using Trimmomatic v0.33, and the trimmed reads were aligned to the GRCm38 reference genome using Hisat2 v2.1.0. The gene-level read counts were generated from the mapped reads using featureCounts. Downstream differential expression analysis was carried out using DESeq2 v3.12 [40] in R v.3.6.2. Multi-dimensional scaling (MDS) plot-diagnostics identified a non-transgenic WT sample as an outlier. Therefore, the sample was excluded from the differential gene expression analysis. A model was built to test the effect of genotype, adjusting for sex (design=∼sex + genotype). Wald test was performed for each gene and the attained p-values were corrected for multiple testing to using the Benjamini-Hochberg method. A differentially expressed gene was defined as FDR < 0.05. Gene set enrichment analysis and the transcription factor-gene co-occurrence analysis were conducted using Enrichr [41, 42] and Ingenuity Pathway Analysis. Volcano plots and Z-score heatmaps were generated in GraphPad Prism 8.

### Statistical Analysis

Data are expressed as means ± standard error (SE). Statistical tests were carried out using R and GraphPad Prism 8. The normality of the data was tested with Shapiro-Wilk test. For comparing multiple groups, Welch’s ANOVA followed by post-hoc Dunnett’s multiple comparison tests were used for normally distributed data, whereas Kruskal-Wallis followed by Wilcoxon rank-sum test was used for non-normally distributed data. For comparing immunoblot results of membrane-associated or phosphorylated proteins between paired samples, the Wilcoxon signed-rank test was used. For comparisons of behavioral performance over consecutive days, repeated measures two-way ANOVA was conducted followed by post-hoc Sidak’s multiple comparisons. To compare differences between two groups, the Student’s t-test was used. The *p*-value < 0.05 was considered statistically significant.

## Results

### Elevated FT expression in AD brains accompanied by an increase in H-Ras farnesylation and activation of downstream signaling

Previous studies have reported upregulation of isoprenoids biosynthesis in AD brains as evidenced by an increase in the level of isoprenoids and in the mRNA expression of farnesyl pyrophosphate synthase (FPPS) and geranylgeranyl pyrophosphate synthase (GGPPS) [23, 24]. However, little is known whether the elevation in isoprenoids leads to increased prenylation of target proteins including small GTPases and activation of their downstream signaling pathways. To evaluate the levels of protein prenylation during the progression of AD, postmortem human brain tissue samples were obtained from the Religious Orders Study (ROS) participants with three different clinical diagnoses –NCI, MCI, and Alzheimer’s dementia [30]. Sixty individuals (20 from each group), matched for age, sex, and education, were selected for this study solely based on clinical diagnosis independent of neuropathological scores. Demographic and pathology information of the selected participants are summarized in **Table 1**.

First, to measure the protein expression of FT and GGT, the brain tissue homogenates were subjected to immunoblot analyses (**Figure 2A**). The results showed that FT levels were significantly elevated in AD brains and a trend of increase in the MCI group (**Figure 2B**), whereas GGT levels were similar among the three groups (**Figure 2C**). Then, we proceeded to determine whether elevated FT leads to an increase in protein farnesylation. Given that prenylation facilitates membrane association of target proteins, we isolated prenylated proteins from unprenylated proteins by separating the membrane and cytosolic fractions of the brain lysates using analytical ultracentrifugation as previously reported [33, 43]. H-Ras, unlike other Ras isoforms such as N-Ras or K-Ras that can be alternatively geranylgeranylated upon inhibition of FT, is exclusively farnesylated [44]. Thus, H-Ras was selected to assess protein farnesylation in the brain samples. Immunoblot analyses showed that membrane-associated (farnesylated) to cytosolic (unfarnesylated) H-Ras ratios were more than 3-fold higher in MCI and 4-fold higher in AD brains compared with NCI (**Figure 2, F** and **G**). Notably, there was no difference in total H-Ras levels in the unfractionated brain tissue homogenates among the three groups (**Figure 2, C** and **D**), indicating the specific increase in the farnesylated H-Ras in MCI and AD brains.

**Figure 2.**
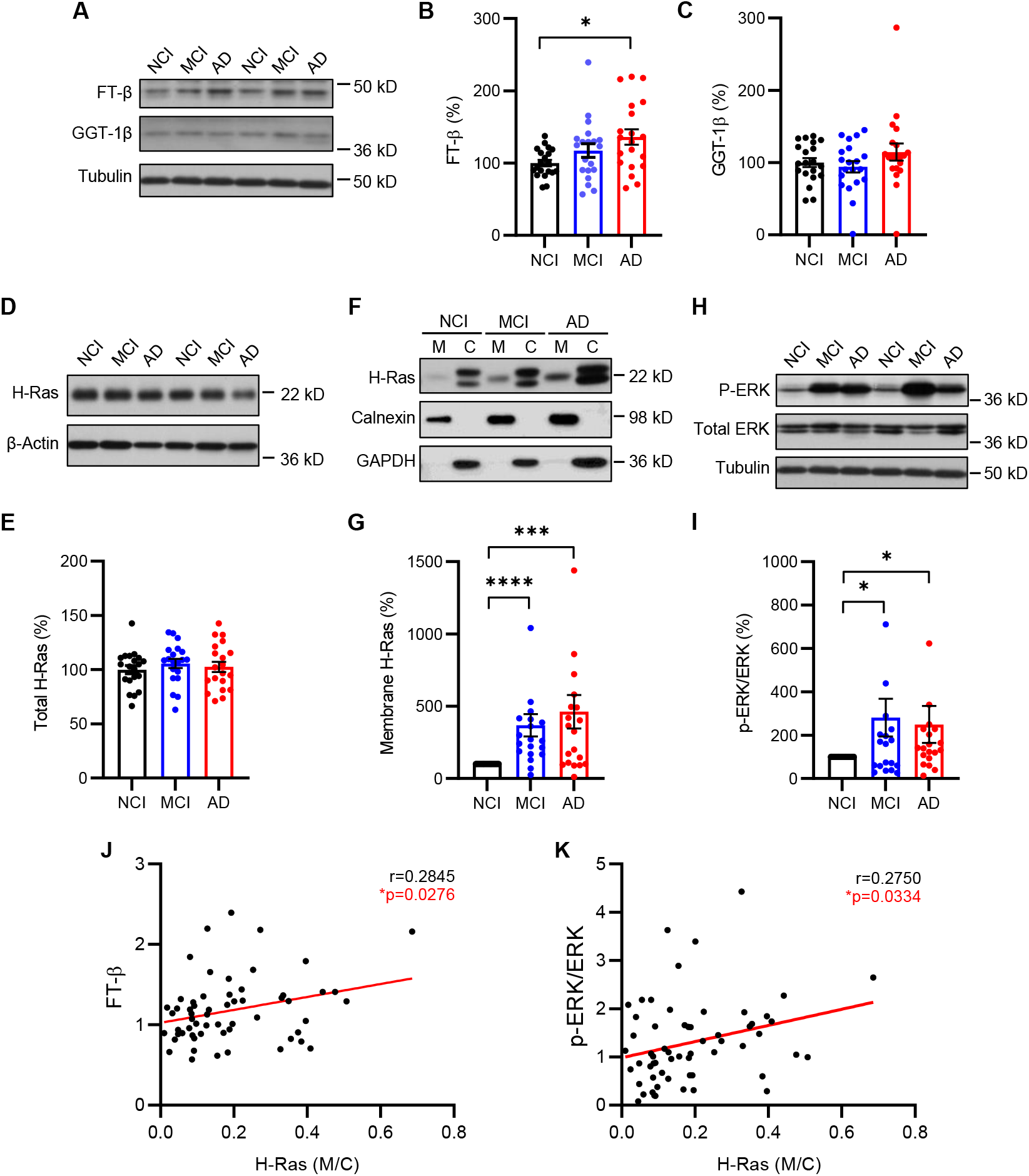
Elevation of FT in AD brains accompanied by an increase in H-Ras farnesylation and ERK activation in brains with MCI as well as AD. (**A**) Immunoblot analysis of FT-β subunit and GGT-1β subunit in the prefrontal cortical tissue lysates from individuals with no cognitive impairment (NCI), mild cognitive impairment (MCI), or Alzheimer’s disease (AD). (**B, C**) Densitometry analysis of FT-β and GGT-1β immunoblots normalized by tubulin with the levels in the NCI group set as 100%. (**D, E**) Immunoblot analysis of total H-Ras and densitometric analysis of H-Ras normalized by β-actin. (**F, G**) Immunoblot analysis of H-Ras in the membrane (M) and cytosolic (C) fractions. Immunoblot analysis of ER membrane-associated calnexin and cytosolic GAPDH confirms the separation of membrane-associated and cytosolic proteins. Ratios of membrane-associated (farnesylated) H-Ras to cytosolic (unfarnesylated) H-Ras were compared relative to the NCI group. (**H, I**) Immunoblot analysis of phospho-ERK (P-ERK) and total ERK in the brain tissue lysates. Ratios of P-ERK (activated) to total ERK were compared relative to the NCI group. (**J, K**) Positive correlations of membrane (farnesylated) H-Ras with FT and P-ERK levels by Pearson correlation analysis. n=20 per group; (B, C and E) Kruskal-Wallis, 2-tailed with post-hoc Dunn’s multiple comparison test; (G and I) Wilcoxon signed rank-order test, 2-tailed; * *p* < 0.05; *** *p* < 0.005; **** *p* < 0.0001.

It is well known that the extracellular signal-regulated kinase (ERK) is a key downstream effector of Ras and its activity is regulated by phosphorylation status [45]. To determine if the elevated farnesylated H-Ras led to changes in the activation of downstream signaling, we measured phosphorylated (activated) and total ERK levels (**Figure 2H**). Consistent with the farnesylated H-Ras levels, phosphorylated-to-total ERK ratios were approximately 2-3 fold higher in MCI and AD brains compared with those with NCI (**Figure 2I**). Further, the levels of membrane/farnesylated H-Ras were positively correlated with the levels of FT (**Figure 2J**) and phosphorylated-to-total ERK ratios (**Figure 2K**), respectively. Taken together, these results suggest that upregulation of FT expression, along with subsequent increase in protein farnesylation and overactivation of downstream signaling pathways, is an early event in the pathogenic cascade of AD.

### Generation of forebrain neuron-specific FT knockout in APP/PS1 mice

Next, we tested whether direct genetic suppression of FT could modify the pathogenic process of AD. Previously, we have shown that germline/systemic FT heterozygous deletion rescues cognitive function and mitigates amyloid pathology as well as neuroinflammation in the APP/PS1 transgenic mouse model of AD [27]. To dissect the specific role of neuronal FT in AD pathogenesis and to circumvent the impact of FT deletion during the embryonic development, we crossed FT-floxed (FT^f/f^) mice with the CaMKIIα promoter-driven Cre recombinase (Cre+) transgenic mice to generate conditional FT knockout mice (FTnKO) (**Figure 3A**), in which FT is selectively deleted in the forebrain neurons postnatally [33, 34]. To confirm the age dependence and brain regional specificity of FT deletion, cortical, hippocampal, and cerebellar tissue lysates of the conditional FT knockout mice were subjected to immunoblot analyses for the β subunit of FT and HDJ-2, a well-known farnesylated protein that is widely used as a marker for farnesylation [46]. As expected, the results showed greater reduction of FT with increasing age, accompanied by an increase in unfarnesylated HDJ-2 in the cortex and hippocampus, but not in the cerebellum (**Figure 3B**). Further, immunoblot analysis of membrane and cytosolic fractions of cortical tissue lysates from FTnKO mice showed reduced membrane localization of FT substrates, H-Ras and Rheb (Ras homolog enriched in brain), while the membrane association of a GGT substrate RhoA were not affected by conditional FT knockout (**Figure 3C**). These results demonstrated the specificity of forebrain FT knockout in these mice. To investigate the role of neuronal FT in the pathogenic process of AD, FT^f/f^ Cre+ mice were crossed with APP/PS1/FT^f/f^ to produce APP/PS1 mice with neuronal FT knockout (APP/PS1/FT^f/f^/Cre+, designated as APP/PS1/FTnKO), and littermate APP/PS1 controls (APP/PS1/FT^f/f^/Cre-, designated as APP/PS1), along with FT^f/f^/Cre+ (FTnKO) and FT^f/f^/Cre-(wild-type, WT) mice.

**Figure 3.**
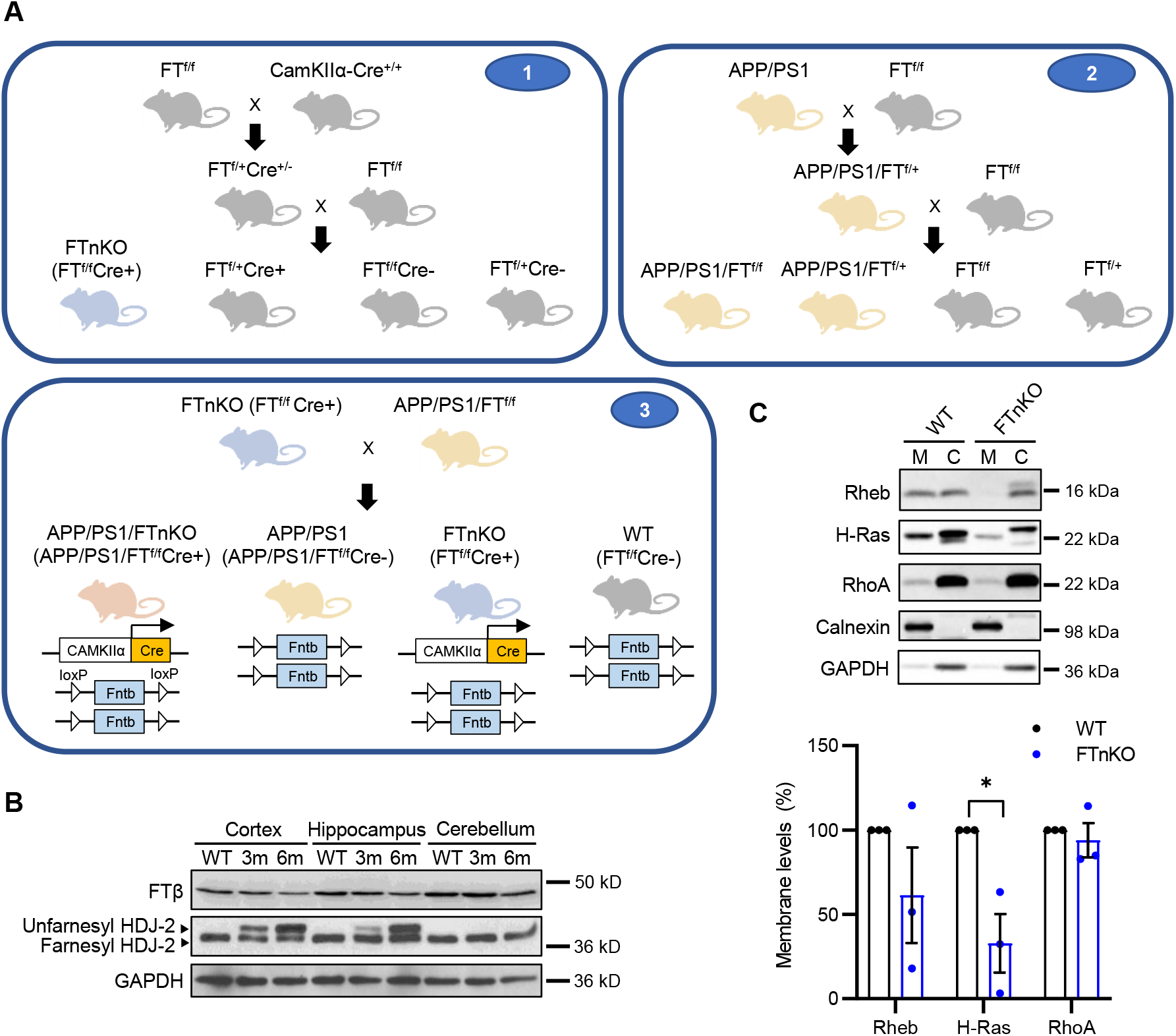
Generation of forebrain neuron-specific FT knockout mice. (**A**) Schematic representation of the 3-step breeding strategy to generate APP/PS1 mice harboring homozygous FT knockout in forebrain neurons (APP/PS1/FTnKO) and littermate controls. (**B**) Representative immunoblot images of FT-β and HDJ-2, a well-known farnesylated protein that is widely used as a marker for farnesylation, in cortical, hippocampal, cerebellar tissue homogenates of WT and FTnKO mice at 3 and 6 months of age. Unfarnesylated HDJ-2 band is detected in the forebrain region (cortex and hippocampus), but not in the cerebellum, of FTnKO mice at 3 months, and the level is elevated at 6 months, confirming both brain region-and age-dependent deletion of FT. (**C**) Immunoblot analysis of representative small GTPases in membrane (M) and cytosolic (C) fractions of cortical tissue lysates of FTnKO mice compared with WT controls (n=3/genotype). Membrane-associated FT substrates, farnesylated H-Ras and Rheb, are reduced significantly or trending reduction in FTnKO compared with WT controls, whereas membrane levels of GGT substrate RhoA are not affected, confirming the specificity of FT deletion. Wilcoxon signed rank-order test, 2-tailed; * *p* < 0.05.

### Improved spatial learning and memory in neuronal FT knockout APP/PS1 mice

To evaluate whether forebrain neuronal FT deletion affects neurobehavioral function, APP/PS1/FTnKO and APP/PS1 mice were subjected to a battery of behavioral tests including the open field test, elevated plus maze test, and the Morris water maze test at 9 months of age. There were no significant differences in locomotor response and exploratory behavior between the two groups assessed by the open field test (**Table 2**). Interestingly, in the elevated plus maze test, APP/PS1/FTnKO mice spent significantly longer time in the open arm than APP/PS1 mice on the second day of the test, suggesting that forebrain neuronal FT deletion reduced anxiety in APP/PS1 mice (**Table 2**). Hippocampus-dependent spatial memory acquisition and retention were evaluated using the Morris water maze test (**Figure 4A**). Even though the mean escape latencies during the 5-day acquisition phase were not significantly different between the two groups, the repeated measures two-way ANOVA revealed a significant interaction between genotype and day during the test, indicating a steeper/faster learning curve in APP/PS1/FTnKO mice (**Figure 4B**). Consistently, the probe trial, conducted 24 hours later to evaluate the memory retention, showed that APP/PS1/FTnKO group spent significantly more time in the target quadrant compared with APP/PS1 group (**Figure 4C**), indicating improved memory performance in APP/PS1/FTnKO mice. To determine whether forebrain neuron-specific FT deletion had any effects on non-cognitive parameters, the visible platform version of the Morris water maze test was conducted. The results showed that there were no significant differences in the escape latencies to reach the visible platform between the two groups (7.7 ± 1.5 sec vs. 6.9 ± 0.8 sec), indicating neuronal FT deletion did not affect the visual acuity or swimming ability. Taken together, these results demonstrated that forebrain neuron-specific deletion of FT reduced anxiety and improved spatial learning and memory function in APP/PS1 mice.

**Table 2.** Exploratory activities and anxiety levels of APPPS1 and APP/PS1/FTnKO mice.

|  | **APPPS1**<br>**(n=13)** | **APP/PS1/FTnKO**<br>**(n=11)** |
| --- | --- | --- |
| **Motor activity (open field)** |  |  |
| Path length (m) |  |  |
| Day 1 | 20.37 ± 4.65 | 13.48 ± 4.59 |
| Day 2 | 14.36±3.88 | 8.95 ± 2.82 |
| Day 3 | 11.92 ± 3.78 | 5.94 ± 1.74 |
| Percentage of time in the central zone (%) |  |  |
| Day 1 | 1.72 ± 0.78 | 1.23 ± 0.74 |
| Day 2 | 0.29 ± 0.14 | 0.05 ± 0.05 |
| Day 3 | 0.28 ± 0.14 | 0.02 ± 0.02 |
| **Anxiety (elevated plus-maze)** |  |  |
| Entries to open arms |  |  |
| Day 1 | 7.53 ± 1.58 | 6.45 ± 1.38 |
| Day 2 | 2.54 ± 0.55 | 7.09 ± 2.01 |
| Percentage of time in open arms |  |  |
| Day 1 | 16.18 ± 3.85 | 37.12 ± 10.51 |
| Day 2 | 6.28 ± 1.93 | 24.08 ± 8.63* |

\* $p < 0.05$ , repeated measures two-way ANOVA with post-hoc Sidak's multiple comparisons.

**Table 3.**
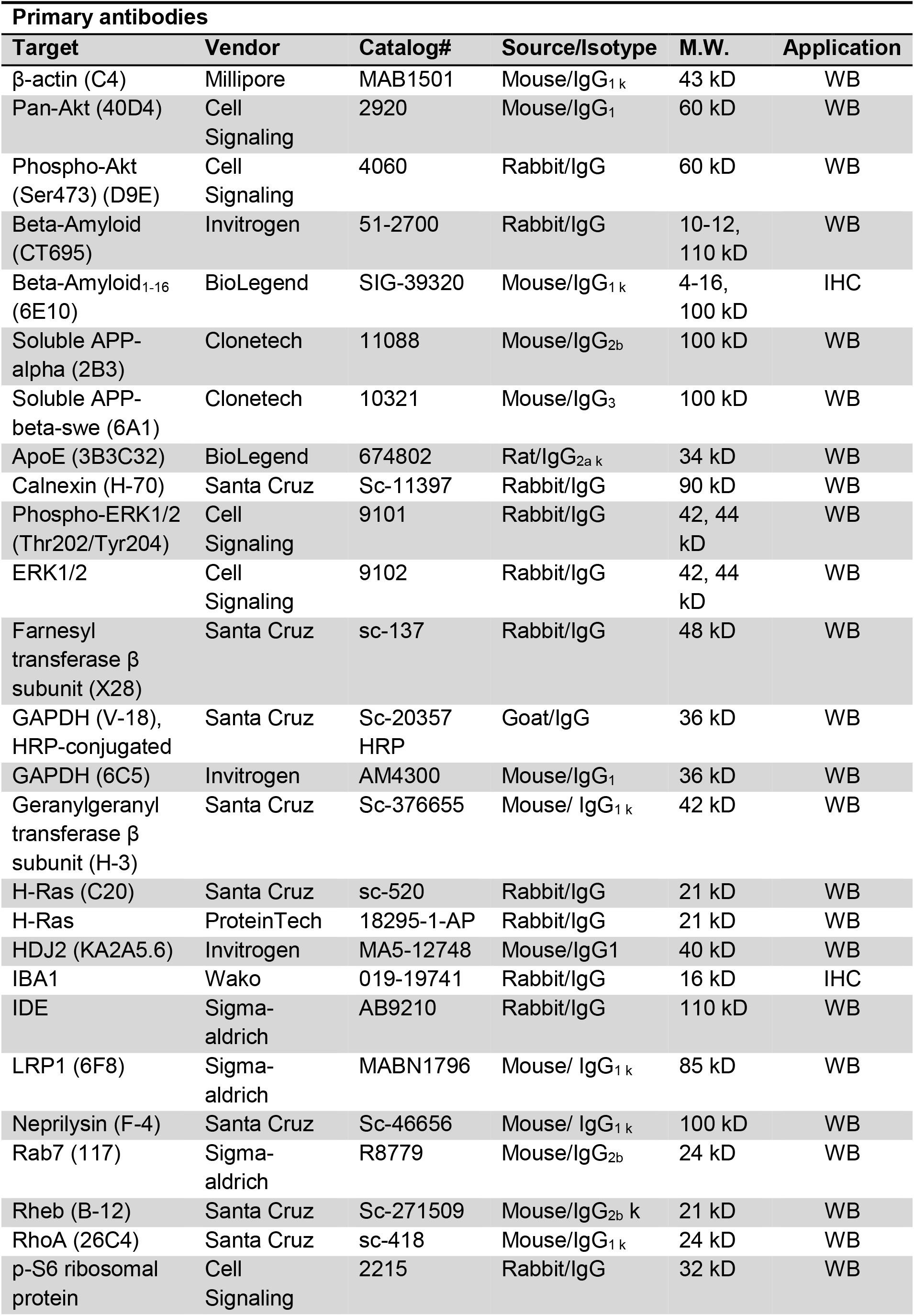
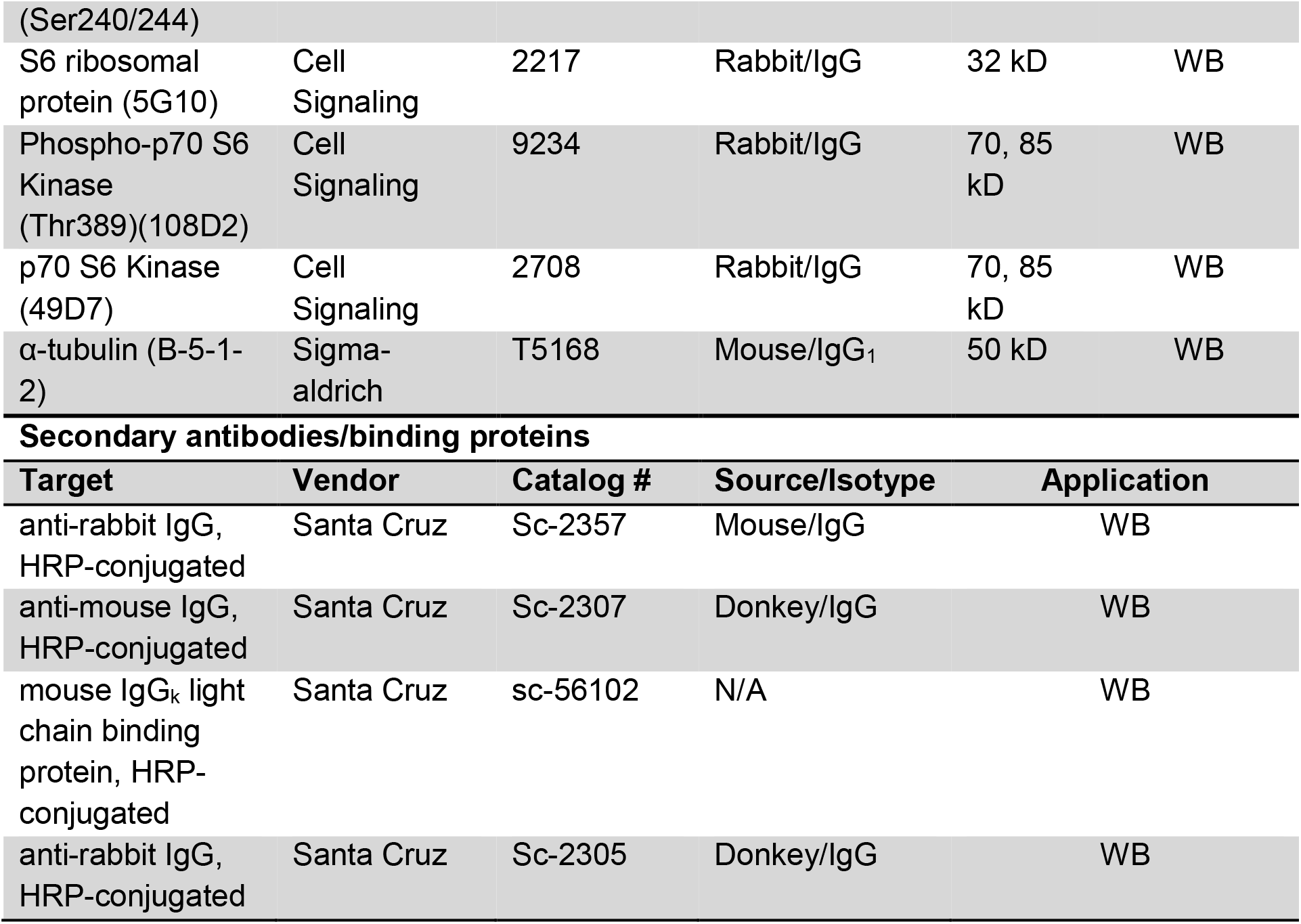
Antibodies used in this study.

**Figure 4.**
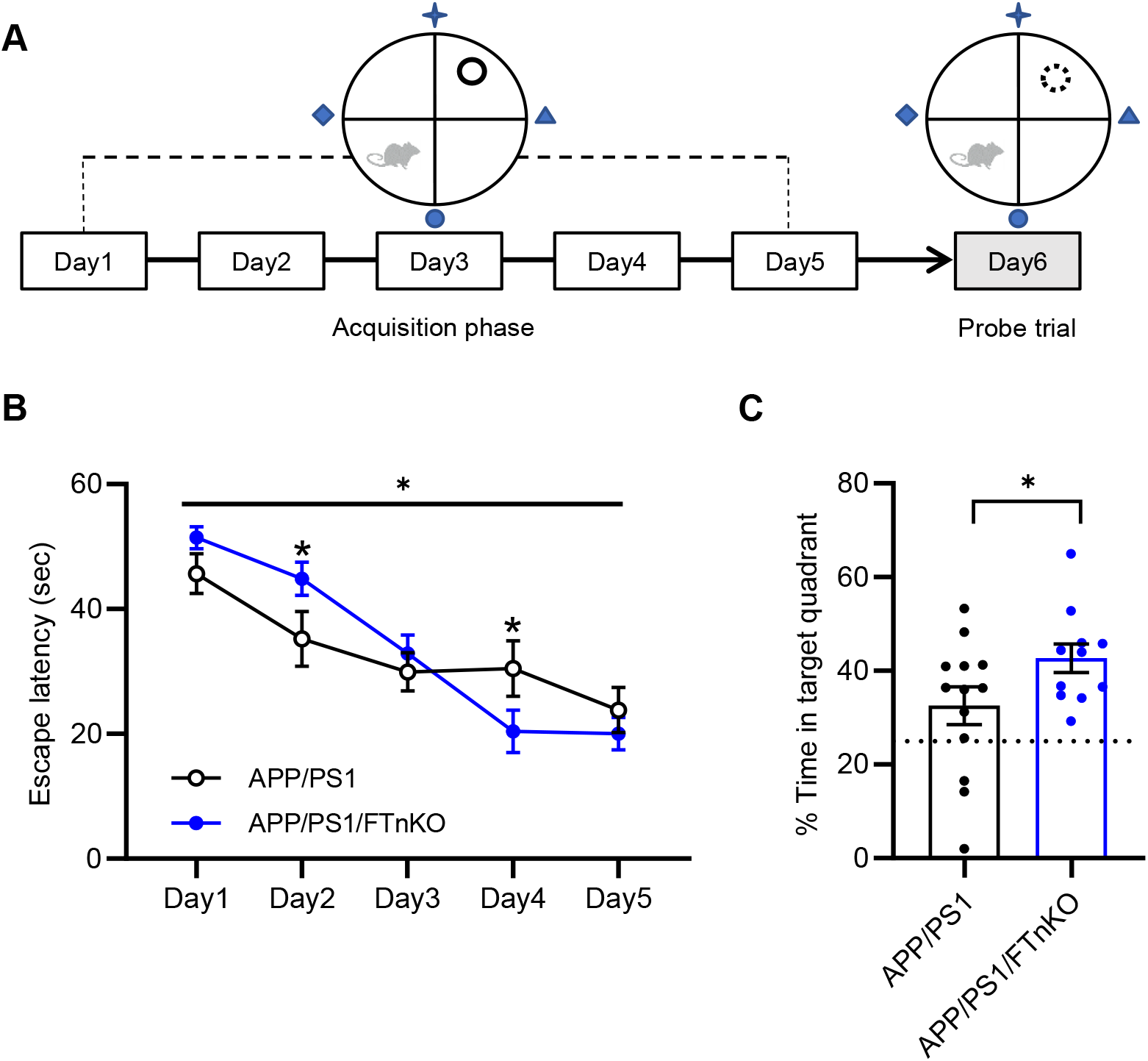
Improvement of spatial learning and memory in neuronal FT knockout APP/PS1 mice. (**A**) Schematic representation of the Morris water maze testing paradigm. (**B**) Escape latencies during the acquisition phase of the test. Repeated measures two-way ANOVA with post-hoc Sidak’s multiple comparisons shows a significant interaction between genotype and day during the test (F(4, 90) = 3.188, *p* = 0.0169), indicating a steeper/faster learning curve in APP/PS1/FTnKO mice; * *p* < 0.05. (**C**) Percentage of time spent in the target quadrant in the probe trial. Time spent in the target quadrant due to chance (25%) is marked with a dashed line. n=11-13 mice/genotype at 9 months; unpaired Student’s t-test, 1-tailed; * *p* < 0.05.

### Reduced cerebral amyloid load and neuroinflammation in neuronal FT knockout APP/PS1 mice

To investigate whether the behavioral improvement in APP/PS1/FTnKO mice was accompanied by any changes in neuropathology, we assessed cerebral amyloid load and associated neuroinflammation. First, we measured Aβ40 and Aβ42 levels in the carbonate-soluble and carbonate-insoluble (guanidine-soluble) fractions in the cortical and hippocampal lysates using Aβ species-specific ELISA. Aβ40 and Aβ42 levels in the carbonate-soluble fraction were markedly reduced by approximately 40-50% in APP/PS1/FTnKO mice compared with APP/PS1 controls in both cortical and hippocampal regions (**Figure 5, A** and **C**). Similarly, both guanidine-soluble Aβ40 and Aβ42 were significantly reduced in the hippocampus. Interestingly, in the cortex, guanidine-soluble Aβ40 was evidently reduced whereas there was only a trend of reduction for Aβ42 (**Figure 5, B** and **D**). Importantly, the total levels of Aβ in both soluble and insoluble fractions of the cortex or hippocampus were substantially reduced in APP/PS1/FTnKO mice compared with APP/PS1 controls (**Figure 5**).

**Figure 5.**
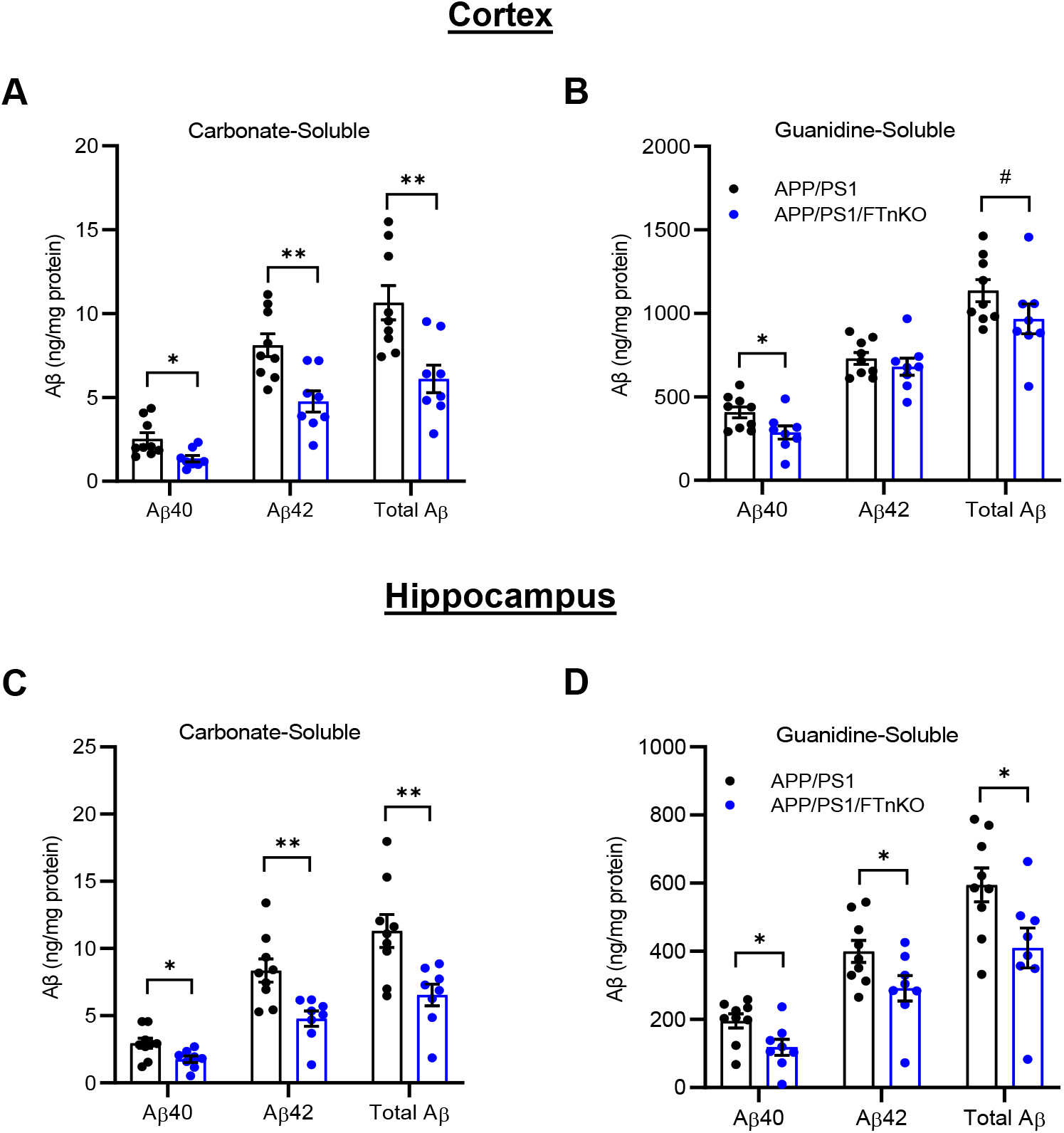
Reduced Aβ accumulation in APP/PS1/FTnKO mice. Aβ was serially extracted from the cortex and hippocampus using carbonate and guanidine buffers. Aβ40 and Aβ42 levels in each fraction were measured by ELISA. (**A)** Carbonate- and (**B**) guanidine-soluble Aβ40 and Aβ42 in the cortex of APP/PS1 and APP/PS1FTnKO mice. (**C**) Carbonate- and (**D**) guanidine-soluble Aβ40 and Aβ42 in the hippocampus of APP/PS1 and APP/PS1FTnKO mice. n=8 mice/genotype; unpaired Student’s t-test, 2-tailed; # *p* = 0.07; * *p* < 0.05; ** *p* < 0.01.

Consistent with the ELISA data, immunostaining of Aβ on brain sections using 6E10 antibody revealed a significant reduction of amyloid plaque deposition in the cortex and hippocampus in APP/PS1/FTnKO mice (**Figure 6, A-C**). Careful inspection of amyloid plaques suggested that forebrain neuron-specific deletion of FT appeared to modify the size of the plaques. Thus, a sub-analysis was performed to quantify the size distribution of 6E10-positive amyloid plaques. Intriguingly, the results showed that in both the cortex and hippocampus, small (<20 µm) and medium (20-50 µm) plaques were significantly reduced, whereas large (>50 µm) plaques were not significantly reduced in APP/PS1/FTnKO mice compared with APP/PS1 mice (**Figure 6, D** and **E**). These results suggest that neuronal FT deletion leads to a substantial reduction in the formation of nascent amyloid plaques, which results in an overall reduction of Aβ deposition in the brain. As β-amyloidosis is often associated with neuroinflammation, the activation status of microglia was assessed next by immunostaining of IBA-1. The results showed a significant reduction in microglial activation in the cortex of APP/PS1/FTnKO compared with APP/PS1, while not in the hippocampus (**Figure 6, G** and **H**). Together, these findings demonstrate that forebrain neuron-specific deletion of FT reduces amyloid pathology and associated neuroinflammation, contributing to the improvement of cognitive function in APP/PS1/FTnKO mice.

**Figure 6.**
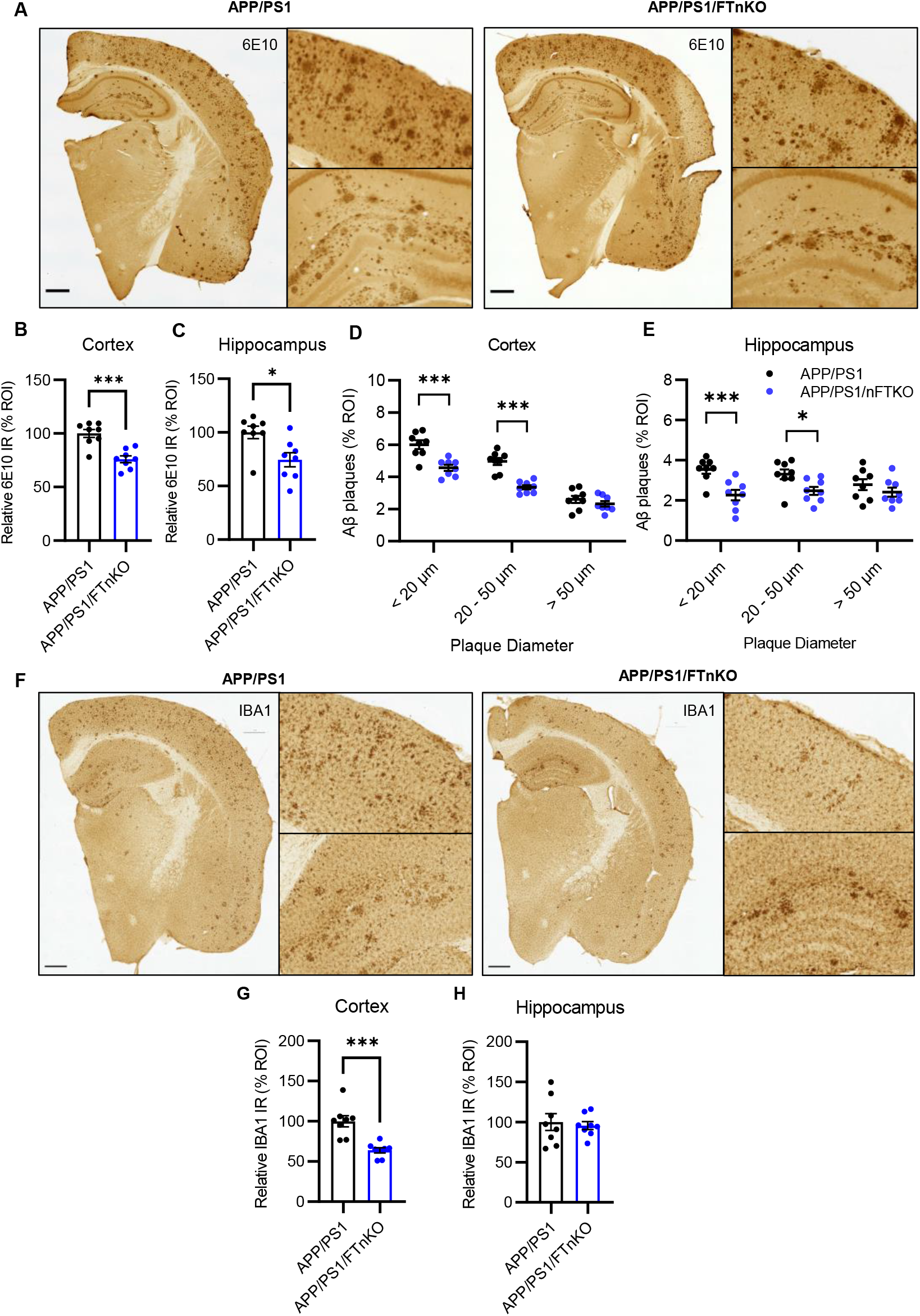
Reduced Aβ deposition in APP/PS1/FTnKO mice. (**A**) Representative brain sections from APP/PS1 and APP/PS1/FTnKO mice immunostained with anti-Aβ antibody (6E10). Scale bar, 500 μm. (**B, C**) Cortex and Hippocampal Aβ loads determined by immunohistochemical and morphometric analyses with the Aβ-positive area in the APP/PS1 control group set as 100%. (**D, E**) Quantification of small (<20 µm), medium (20-50 µm), and large (>50 µm) amyloid plaques in the cortex and hippocampus. **(F**) Representative brain sections from APP/PS1 and APP/PS1/FTnKO mice immunostained for activated microglia using anti-IBA1 antibody. Scale bar, 500 μm. (**G, H**) Microglial activation in the cortex and hippocampus determined by morphometric analyses with the IBA1-positive area in the APP/PS1 control group set as 100%. n=8 mice/genotype; Student’s t-test, 2-tailed; * *p* < 0.05; *** *p* < 0.001.

### Reduced proteolytic processing of APP in neuronal FT knockout APP/PS1 mice

To elucidate the mechanisms underlying the reduction of Aβ load in APP/PS1/FTnKO mice, we studied both Aβ generation and clearance pathways. To assess the proteolytic processing of APP, we measured the steady-state levels of full-length APP, soluble amino-terminal fragments of APP (sAPP), and carboxyl-terminal fragments (CTF) produced by α- and β-secretase cleavages in the brain lysates of APP/PS1/FTnKO and APP/PS1 mice. Immunoblot analysis showed that neuron-specific FT deletion did not change the expression of full-length APP but significantly reduced the production of sAPPα and CTFβ, with a trend reduction in sAPPβ and CTFα (**Figure 7A**). Importantly, there was a trend reduction in the CTFβ/α ratio and the total CTF level was significantly reduced in the brain lysates of APP/PS1/FTnKO mice compared with APP/PS1 mice (**Figure 7B**). These results indicate that neuron-specific FT deletion reduces overall APP processing in neurons; more so in the amyloidogenic processing by β-secretase, without affecting the total levels of APP, thereby leading to subsequent decreases in Aβ production.

**Figure 7.**
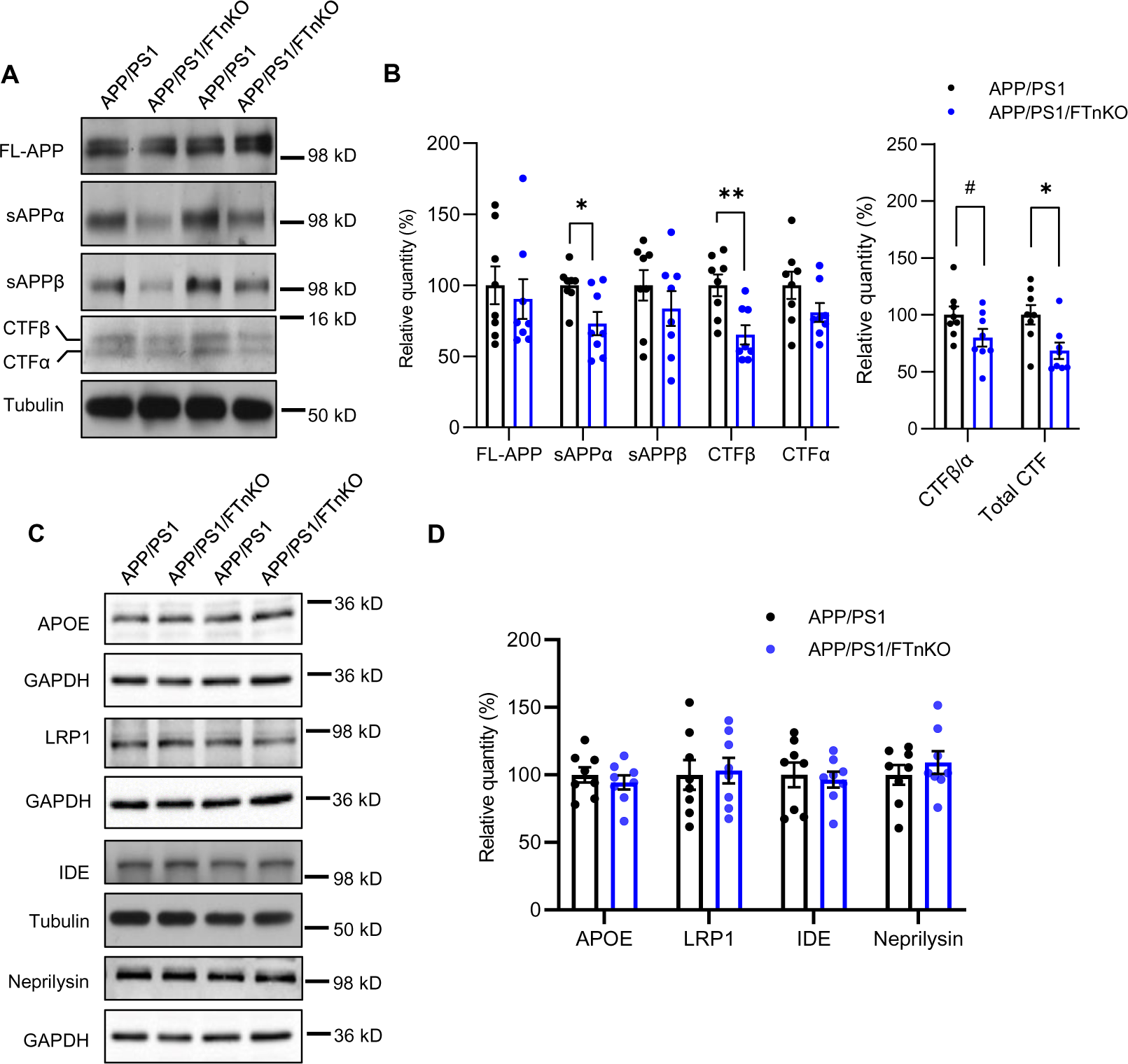
Reduced APP proteolytic processing in APP/PS1/FTnKO mice. (**A, B**) Immunoblots and densitometric analysis of full-length APP (FL-APP), and APP cleavage products, carboxyl-terminal fragments (CTFα and CTFβ), and amino-terminal fragment of APP (sAPPα and sAPPβ). Protein levels were normalized by the amount of tubulin with the levels in the APP/PS1 control group set as 100%. (**C, D**) Immunoblots and densitometric analysis of proteins involved in Aβ clearance pathways including APOE, LRP1, IDE, and neprilysin. Protein levels were normalized by the amount of tubulin or GAPDH, and the levels in the control group set as 100%. n=8/genotype; Student’s t-test, 2-tailed; # *p* = 0.08; * *p* < 0.05; ** *p* < 0.01

Next, we assessed whether forebrain neuronal FT deletion affected Aβ clearance pathways. Prior studies have established that APOE and its receptor LRP1 [47] as well as Aβ-degrading enzymes, including insulin degrading enzyme (IDE) and neprilysin [48], play key roles in Aβ clearance in the brain. Thus, the steady-state levels of APOE, LRP1, IDE, and neprilysin were measured by immunoblot analysis. The results showed no significant differences in the levels of these proteins between APP/PS1/FTnKO and APP/PS1 mice (**Figure 7, C** and **D**), indicating that neuronal FT deletion does not affect these Aβ clearance or degradation pathways.

### Neuronal FT deletion counteracts aberrant mTORC1 activation and downstream lipogenesis in APP/PS1 mice

To gain more insights into the global impact of neuronal FT deletion on gene expression in the brain and the potential molecular mechanisms underlying the beneficial effects of neuronal FT suppression, we performed unbiased transcriptomic analysis on the frontal cortical samples from a cohort of APP/PS1/FTnKO, APP/PS1, and non-transgenic WT controls. There were 156 differentially expressed genes (DEGs) identified in APP/PS1/FTnKO compared with APP/PS1. Of these, 72 genes were significantly downregulated, and 84 genes were upregulated (**Figure 8A**). Functional enrichment analysis of DEGs using MSigDB (Molecular Signatures Database v5.2, http://software.broadinstitute.org/) Hallmark Gene set identified cholesterol homeostasis as the most significantly downregulated pathway followed by the mammalian target of rapamycin complex 1 (mTORC1) signaling pathway (**Figure 8B** and **Figure 9B**).

**Figure 8.**
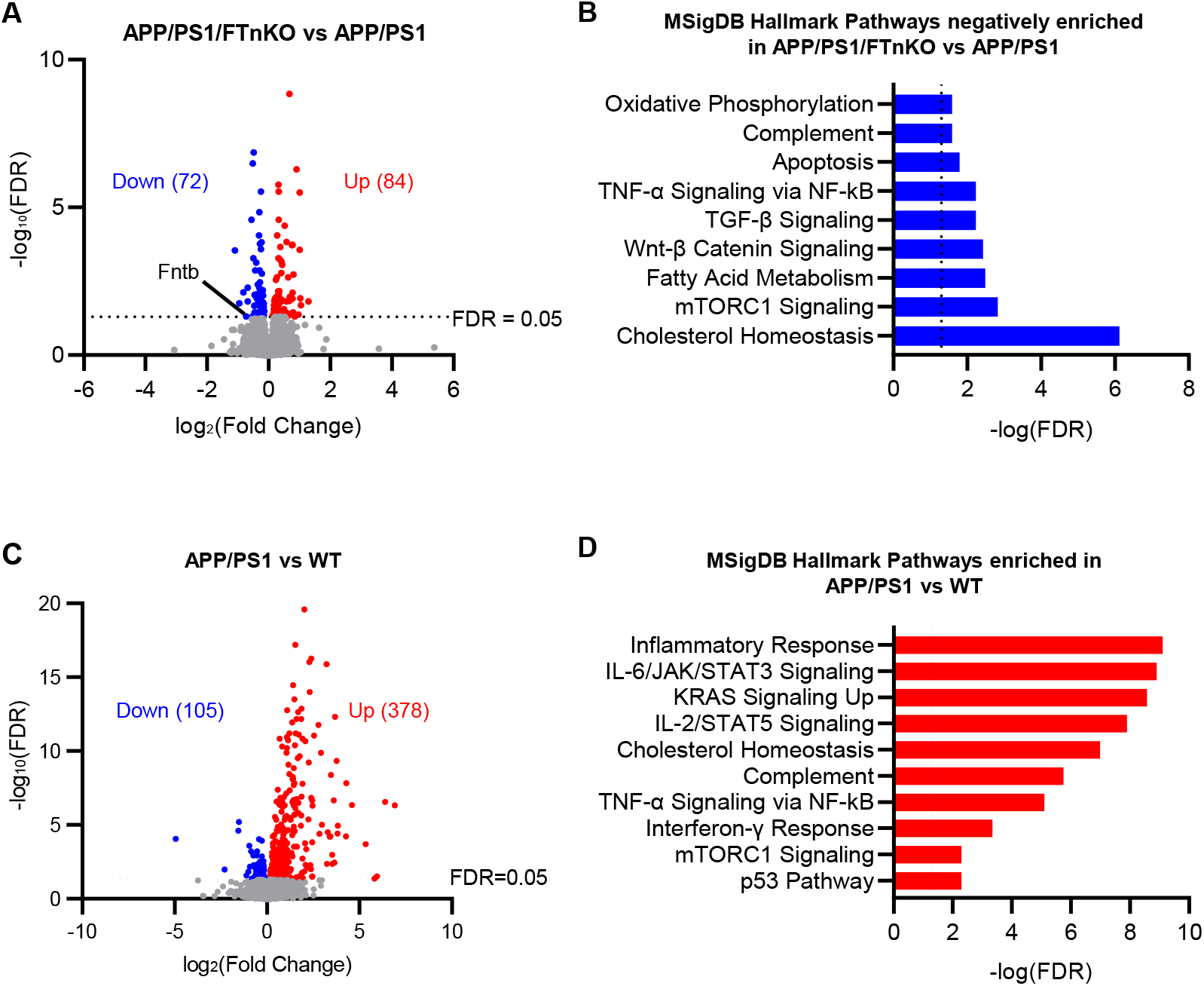
Transcriptomic analysis of frontal cortex from APP/PS1/FTnKO, APP/PS1, and WT mice. (**A**) Volcano plot for cortical transcriptome analysis of APP/PS1/FTnKO versus APP/PS1 mice at 9-10 months (n=6/genotype). Red and blue points represent upregulated and downregulated genes below the significance threshold (FDR < 0.05), respectively. (**B**) Top enriched pathways categorized by mSigDB Hallmark gene set database in differentially downregulated genes in APP/PS1/FTnKO mice compared with APP/PS1 controls. (**C**) Volcano plot for cortical transcriptome of WT (n=5) versus APP/PS1 mice (n=6) at 9-10 months. (**D**) Top enriched mSigDB Hallmark pathways of significantly upregulated genes (FDR < 0.05) in the cortex of APP/PS1 mice compared with WT controls.

**Figure 9.**
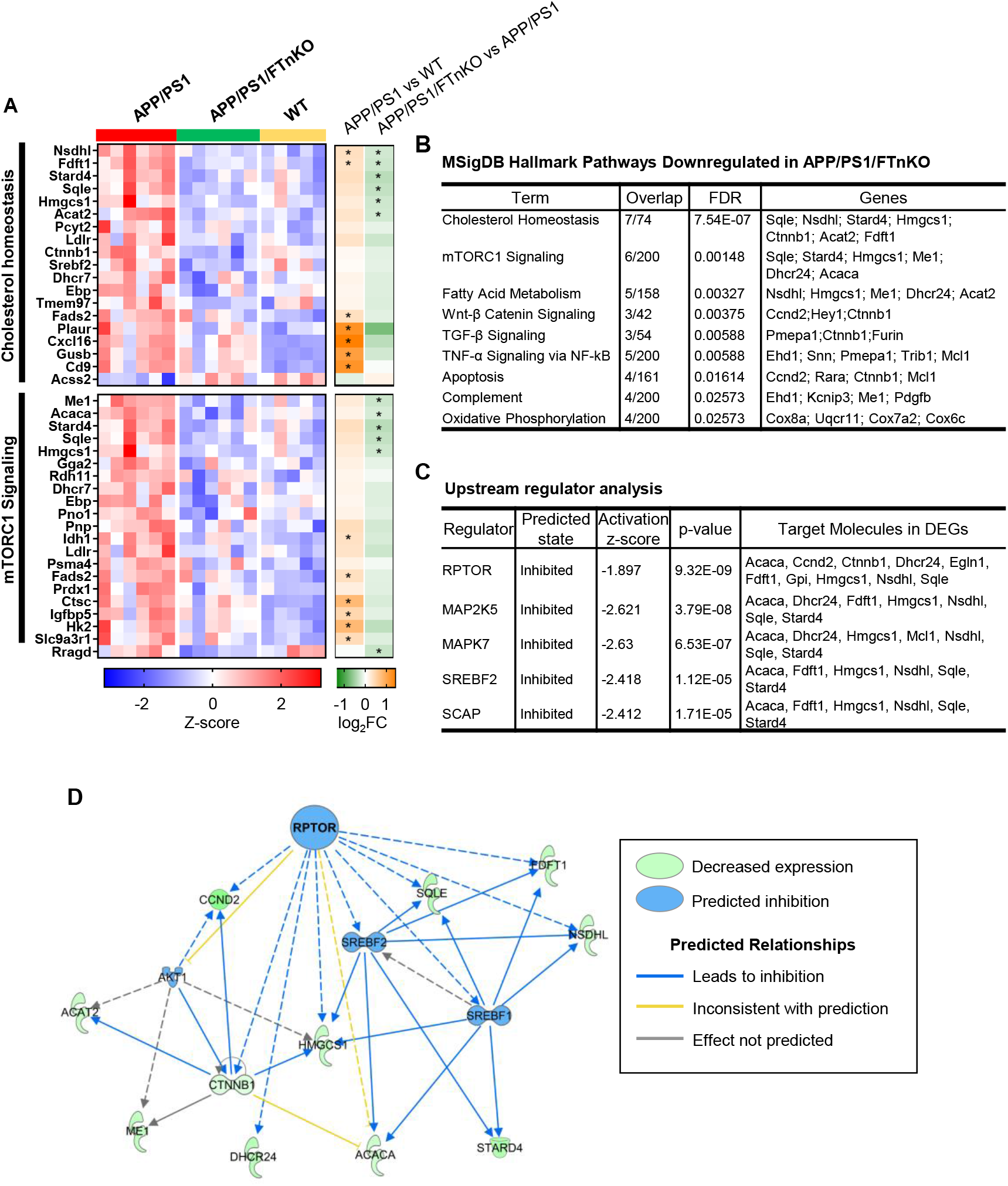
Normalization of mTORC1 and SREBP-mediated lipogenesis pathways in the cortex of APP/PS1/FTnKO mice. (**A**) Heatmap of RNA expression z-scores computed for selected genes in the cholesterol homeostasis and mTORC1 signaling pathways across the three genotypes. The color gradient of the bars on the right reflect the log2(fold change) of the genes, and the genes that are significantly downregulated or upregulated (FDR < 0.05) are indicated by an asterisk (*). (**B**) Table of top downregulated mSigDB Hallmark pathways and the corresponding differentially expressed genes (DEGs) in APP/PS1/FTnKO mice compared with APP/PS1 mice. (**C**) Table of key transcriptional regulators identified through upstream regulator analysis using Ingenuity Pathway Analysis (IPA) software. Based on the log2(fold change) of downstream target genes, the activity of the upstream regulator is predicted using activation z-score. (**D**) Network of Raptor, a regulatory associated protein of mTOR, and its targeted genes. Edges (lines between nodes) represent direct (solid lines) and indirect (dashed lines) interactions as supported by information in the Ingenuity knowledge base.

Pertinently, functional enrichment analysis of 483 DEGs in APP/PS1 mice compared with non-transgenic WT controls indicated an aberrant upregulation of genes that are involved in cholesterol homeostasis and mTORC1 signaling in APP/PS1 mice (**Figure 8, C** and **D**). Z-score heatmap of cholesterol homeostasis and mTORC1 signaling gene clusters shows increased expression of selected genes within these two functional clusters in APP/PS1 mice compared with WT, and notably, neuronal FT deletion reduced the expression levels of many genes within these clusters to comparable levels as in WT mice (**Figure 9A**). mTORC1 is known to activate the sterol regulatory element binding proteins (SREBPs), transcription factors that regulate the expression of genes involved in lipogenesis by promoting the release of SREBPs from the SREBP cleavage-activating protein (SCAP) in the endoplasmic reticulum (ER) and translocation of SREBPs to the Golgi [49, 50]. In the Golgi, SREBPs undergo proteolytic cleavages by site-1 and site-2 proteases, liberating the soluble N-terminal mature SREBPs, which then translocate to the nucleus and bind DNA, upregulating the expression of genes for lipid/sterol synthesis. Therefore, reduced mTORC1 signaling is likely driving the downregulation of cholesterol/lipogenic genes in APP/PS1/FTnKO mice, normalizing them to the levels as in WT mice (**Figure 9, A** and **B**).

To examine the potential upstream regulators that drive the transcriptome changes in APP/PS1/FTnKO mice, we performed an upstream regulator analysis using Ingenuity Pathway Analysis (IPA) software. As expected, the inhibition of Raptor/mTORC1 was shown as the top key upstream regulator of observed transcriptome changes with neuronal FT deletion, followed by the inhibition of downstream SREBP2 and SCAP (**Figure 9, C** and **D**).

### Pharmacological inhibition of Ras and Rheb farnesylation reverses aberrant activation of downstream mTORC1 signaling in APP-overexpressing SH-SY5Y cells

To corroborate the findings from transcriptomic analyses, we tested whether overexpression of APP, thereby increasing Aβ production, leads to overactivation of Ras and Rheb downstream signaling in neuronal cells. We compared the phosphorylation status of ERK and S6 ribosomal protein, surrogate indicators of Ras and Rheb activation, in stably transfected APP695-overexpressing SH-SY5Y (SH-SY5Y-APP695) cells with that in mock-transfected control SH-SY5Y cells. Immunoblot analysis shows that phosphorylated ERK and S6 are significantly elevated in SH-SY5Y-APP695 cells, confirming abnormal upregulation of Ras-ERK and Rheb-mTOR-S6 pathways in neuronal cells overexpressing APP (**Figure 10, A** and **B**).

**Figure 10.**
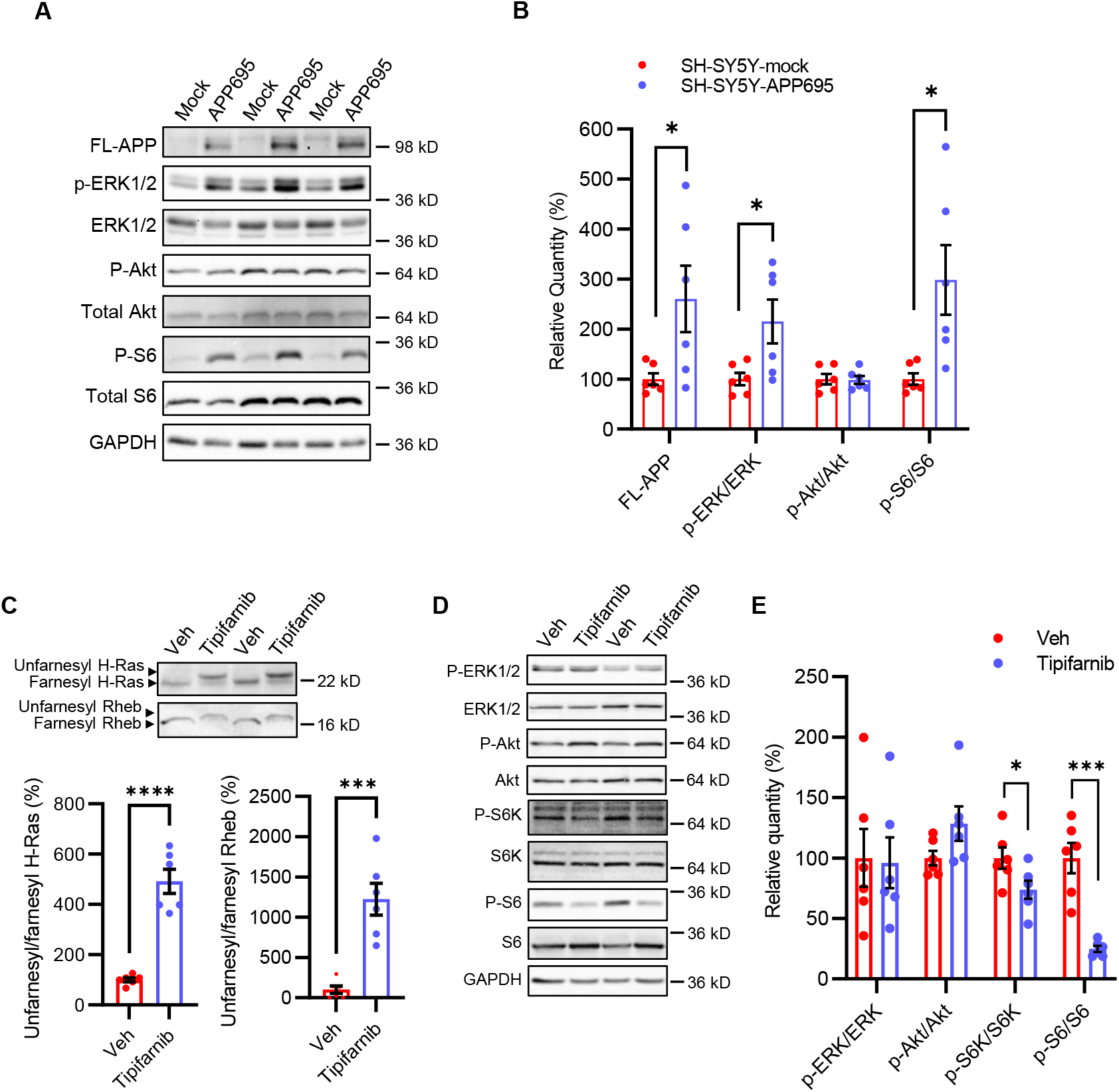
Pharmacological inhibition of FT reverses the hyperactivation of Ras-Rheb-mTORC1 signaling in SH-SY5Y-APP695 cells. (**A**) Immunoblot analysis of FL-APP and downstream targets of Ras-Rheb-mTORC1 signaling, including ERK and ribosomal protein S6, in the lysates of SH-SY5Y-APP695 mock-transfected control cells. (**B**) Densitometric analysis of immunoblots normalized by the amount of GAPDH or total proteins for phosphorylated proteins (P-ERK, P-Akt, P-S6) with the levels in the control group set as 100%. n=6 replicates from 2 independent experiments. (**C**) Immunoblot analysis of farnesylation status of H-Ras and Rheb, and (**D**) phosphorylation status of downstream targets of Ras and Rheb in the lysates of SH-SY5Y-APP695 WT cells after treated with vehicle (DMSO) or 1 μM tipifarnib for 48 hours. (**E**) Densitometric analysis of immunoblots of P-ERK, P-S6K, and P-S6 normalized by the amount of total ERK, S6K, and S6 with the levels in the control group set as 100%. n=6 replicates from 2 independent experiments. Student’s t-test, 2-tailed; * *p* < 0.05; *** *p* < 0.001; **** *p* < 0.0001.

Next, we sought to answer whether pharmacological inhibition of FT using a specific inhibitor tipifarnib can negate the aberrant hyperactivation of Ras and Rheb-mTORC1 signaling pathways in SH-SY5Y-APP695 cells. After 48 hours of tipifarnib treatment in growth media, farnesylation status of H-Ras and Rheb as well as phosphorylation status of ERK, p70 S6 kinase (S6K), and S6 were measured using immunoblots. As expected, we observed clear mobility shift of H-Ras and Rheb bands as most of these proteins became unfarnesylated in the presence of tipifarnib (**Figure 10C**). Moreover, tipifarnib treatment also led to a significant reduction in phosphorylated S6K and S6 (**Figure 10D**), indicating that neuronal FT inhibition reverses amyloid-associated mTORC1 activation. Interestingly, tipifarnib did not affect the ERK phosphorylation even though most of H-Ras became unfarnesylated after 48 hours of treatment (**Figure 10, C** and **D**). This may be explained by the fact that ERK is also phosphorylated by other Ras isoforms such as N-Ras and K-Ras that can be prenylated by GGT when FT is inhibited. Taken together, these results show that inhibition of protein farnesylation reverses aberrant activation of Ras and Rheb-mTORC1 signaling associated with APP/Aβ overproduction.

## Discussion

Findings from this study support the growing evidence that protein prenylation, particularly farnesylation, regulates the pathogenic process of AD. Previously, we have shown that systemic FT haplodeficiency rescues cognitive function as well as reducing Aβ load and neuroinflammation in APP/PS1 mice [27]. In the present study, we found that FT is upregulated in the brain of individuals with AD. Further, membrane-associated H-Ras, an exclusively farnesylated protein, and its downstream effector ERK activation are elevated in individuals with MCI as well as AD, suggesting upregulation of protein farnesylation and its downstream signaling pathways is an early event with primary importance in the pathogenic cascade of AD. Moreover, we show that suppressing forebrain neuronal FT alone leads to improved cognitive function and reduced neuropathology in APP/PS1 mice, revealing the specific role of neuronal FT in the pathogenesis of AD. In addition, transcriptomic analysis uncovers the global impact of neuron-specific FT deletion on cerebral gene expression and identifies the reversal of pathogenic overactivation of mTORC1 signaling network as the top contributor to the beneficial effects in APP/PS1/FTnKO mice, which is also supported by *in vitro* studies with APP-overexpressing neuronal cells.

Prior work with human brain tissue samples has shown an elevation of isoprenoid biosynthesis in AD brains compared with controls, reflected by an increase in the level of FPP and GGPP and the mRNA expression of FPPS and GGPPS [23, 24]. However, whether the protein prenylation process *per se* is affected in AD is not known. Here, studying the brain tissue samples from individuals at different stages of the disease, we found that FT is abnormally upregulated in AD, leading to subsequent elevations of protein farnesylation and overactivation of downstream signaling pathways, as evidenced by an increase in membrane-associated H-Ras and its downstream ERK phosphorylation. Notably, the elevation of membrane-associated H-Ras and ERK phosphorylation occurs in the brain with MCI as well as AD, indicating dysregulation of protein farnesylation and downstream signaling in the early stage of AD. Consistently, other studies have shown that the level of Ras (both cytosolic and membrane/prenylated fractions) in the brain, although without the identification of specific Ras isoforms, is increased prior to both amyloid and neurofibrillary tangle pathologies [51, 52]. Increased expression of Ras and activation of downstream signaling correlate with Aβ levels in AD brains [19]. Importantly, here we show a particular increase in the membrane-associated H-Ras, an exclusively farnesylated protein, in both MCI and AD brains, highlighting the significance of protein farnesylation in the pathogenesis of AD.

Studies in cellular and animal models corroborate the role of protein farnesylation in multiple aspects of AD. Pharmacological treatment with farnesyltransferase inhibitors (FTI) has been shown to modulate synaptic and cognitive function as well as amyloid and tau pathologies [18-20, 53], although potential off-target effects of pharmacological agents may confound the results. Targeting FT directly by the genetic approach, we have reported that systemic FT haplodeficiency rescues cognitive function in addition to reducing amyloid pathology in APP/PS1 mice (27). Here we show that postnatal deletion of neuron-specific FT deletion was sufficient to ameliorate memory deficit and reduce Aβ load in these mice. Interestingly, we also found that neuronal FT deletion significantly reduced anxiety levels in APP/PS1 mice, which was not observed in systemic FT haplodeficient mice. This finding is similar to anxiolytic-like behavior changes reported in rats chronically treated with statins, potentially by modulating N-methyl-D-aspartate (NMDA) receptors in brain regions [54, 55]. Along with the evidence that statin treatment enhances NMDA receptor-mediated synaptic function by inhibiting FPP production and farnesylation (18, 53), these findings suggest that farnesylation-dependent mechanisms in neurons underlie the effects of statins on cognitive function and anxiety levels.

As in systemic FT haplodeficient mice, neuron-specific FT deletion also led to reduced Aβ load in the brain of APP/PS1 mice, indicating the major contribution of neuronal FT-mediated lowering of Aβ to the attenuation of cerebral amyloid pathology. Similarly, the underlying mechanisms involved the shift of amyloidogenic to non-amyloidogenic processing of APP, although neuron-specific FT deletion resulted in a greater reduction of overall process of APP by both α- and β-secretases than systemic FT haplodeficiency [27]. Furthermore, IDE, one of the major Aβ degrading enzymes, was upregulated in the brain of systemic FT haplodeficient mice, contributing to the reduction of Aβ pathology [27], whereas the level of IDE was not changed in the brain of APP/PS1/FTnKO mice. This finding is consistent with the fact that IDE expression is relatively low in neurons compared with glial cells [56], thus the impact of neuronal FT deletion is not sufficient to influence the overall level of IDE in the brain. In agreement with the observation in systemic FT haplodeficient mice, FTI treatment induces IDE secretion from microglia and increases the degradation of Aβ [57]. Interestingly, neuronal FT deletion, like systemic FT haplodeficient, also led to attenuation in microglial activation in the brain, apparently due to reduced amyloid deposition. Taken together, these results demonstrate that while both APP processing and Aβ degradation pathways contribute to reduced cerebral Aβ load in systemic FT haplodeficient mice, significant diminution of overall APP processing is the primary mechanism by which neuronal FT deletion modifies Aβ pathology.

Further molecular insights were provided by transcriptome analysis of frontal cortical tissue samples from different genotypes of mice. Compared with non-transgenic WT controls, the aberrant upregulation of mTORC1 signaling and cholesterol synthesis/metabolism was identified in APP/PS1 mice. Activation of mTORC1 is regulated by various growth factors and oxidative stress [58], which promotes anabolic processes including lipogenesis through SREBP activation and protein synthesis, while inhibits autophagy [59-61]. These findings are consistent with previous studies that have shown an aberrant hyperactivation of mTORC1 pathways along with the accumulation of amyloid plaques in various transgenic AD mouse models [62-64] as well as in postmortem human AD brains [65, 66]. Moreover, genetic reduction or early pharmacological suppression of mTOR has been shown to rescue cognitive deficits, reduce Aβ deposition and intracellular tau accumulation along with an increase of autophagy induction in AD mice [64, 67, 68]. Activation of mTORC1 also stimulates cholesterol/fatty acid synthesis through activation of SREBP1/2 [49, 50]. Our differential gene expression analysis indicates that neuronal FT deletion significantly reduces mTORC1 signaling, and the downstream SREBP1/2-mediated lipogenesis in APP/PS1 mice, restoring them to normal levels as in WT mice.

The reduction of mTORC1 signaling is most likely mediated through the reduction of farnesylated Ras and Rheb in APP/PS1/FTnKO mice, which is supported by results from *in vitro* study with APP-overexpressing neuronal cells. Farnesylation of Ras is required for its localization to the plasma membrane localization, where it interacts with membrane surface receptor tyrosine kinases (RTKs) and Ras guanidine exchange factor (GEF) Sos1 [69]. Ras mediates the activation of RTKs signaling through downstream Raf-MEK-ERK and PI3K-PDK1-Akt signaling cascades, eventually leading to the inactivation of TSC1/2 complex via phosphorylation [70]. TSC1/2 is a GTPase-activating protein that hydrolyzes the active Rheb-GTP to the inactive Rheb-GDP. The inactivation of TSC1/2 through Ras-mediated RTKs signaling increases the active GTP-bound Rheb, which then directly activates mTORC1 [70, 71]. Farnesylation of Rheb is essential for the lysosomal membrane-localization of Rheb, where it encounters and activates mTORC1 [72-74]. Notably, Rheb is a major protein identified by our recent prenylomic analysis, the farnesylation of which is significantly reduced in the brain of FTnKO mice [29], as well as shown in this study by immunoblot analysis of membrane and cytosolic fractions of brain homogenates. Thus, these results support the notion that neuronal FT deletion suppresses the pathogenic mTORC1 overactivation by reducing RTKs-Ras mediated Rheb activation and Rheb-mTORC1 interaction through inhibition of farnesylation in APP/PS1 mice.

Suppression of mTORC1 signaling leads to downregulation of SREBP1/2-mediated cholesterol/lipogenic pathways, which may explain some beneficial outcomes in APP/PS1/FTnKO mice. The role of cholesterol in AD has been extensively studied. In particular, increase of cholesterol in cell membrane/lipid raft microdomains diminishes membrane fluidity, favoring amyloidogenic APP processing by β-secretase in lipid-rafts while reducing neuroprotective, non-amyloidogenic APP processing in non-lipid rafts of the plasma membrane [75-77]. Consistently, modulation of cellular cholesterol influences Aβ production by affecting the APP interactions with β- or γ-secretase [78-80]. In addition, overexpression of SREBP2 in APP/PS1 transgenic mice has been shown to accelerate Aβ accumulation and induce tau tangle formation [81]. Here we show that neuronal FT deletion corrects the heightened SREBP1/2-mediated cholesterol/lipogenic pathways in APP/PS1 mice, thus promoting non-amyloidogenic processing of APP and lowering Aβ production.

In summary, the present study unveils the dynamics of protein farnesylation in the pathogenic process of human AD and provides novel insights into the specific impact of neuronal FT on AD-like memory impairment and amyloid pathology in APP/PS1 mice. Our results demonstrate that abnormal upregulation of FT occurs in the early stage of human AD, and suppressing neuronal FT alone confers neuroprotective benefits against cognitive deficits and amyloid pathology through reversing pathogenic hyperactivation of mTORC1 signaling in APP/PS1 mice (**Figure 11**). In light of the recent approval of an FT inhibitor for the treatment of progeria [82], clinical trials investigating the therapeutic potential of this class of drugs for AD are warranted.

**Figure 11.**
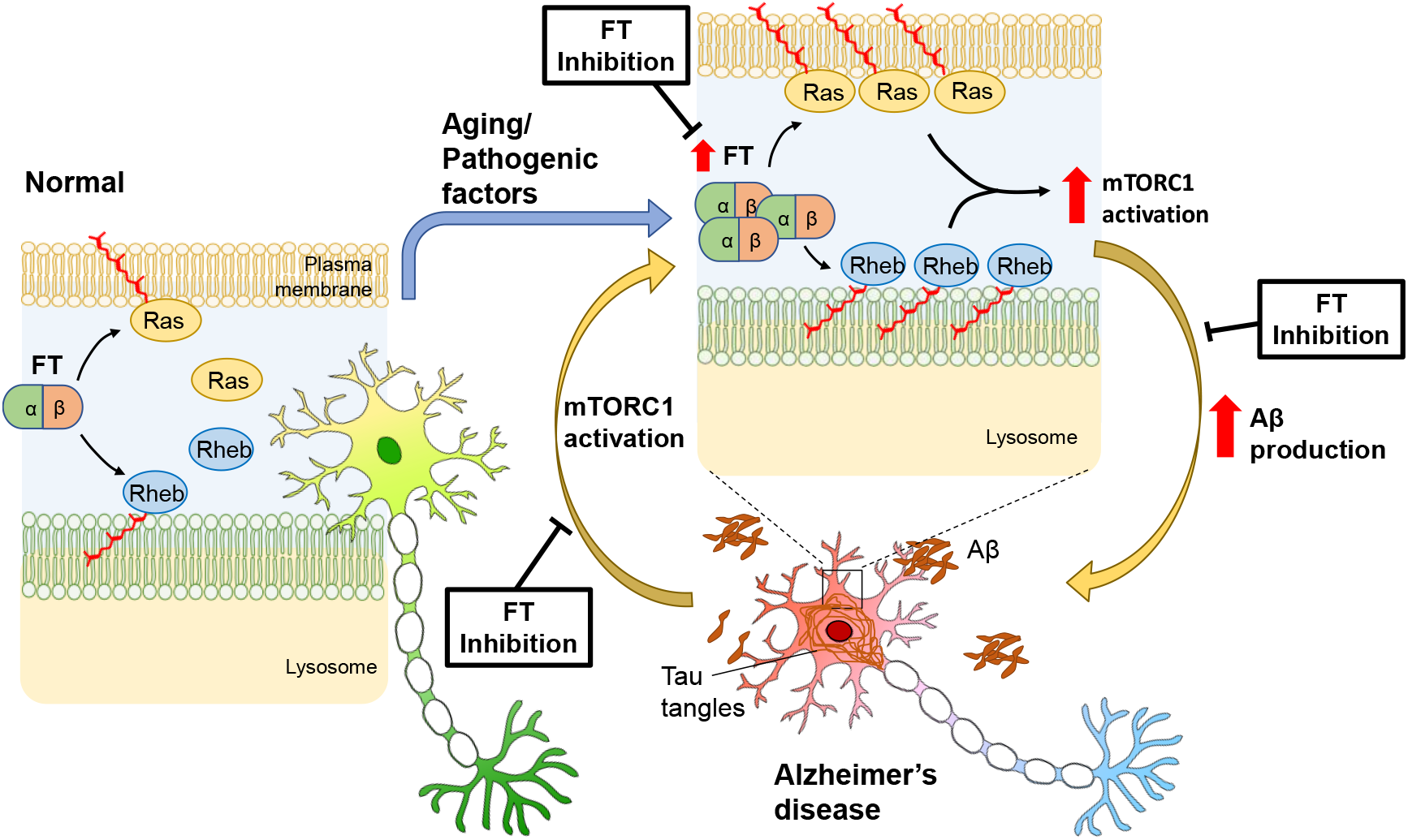
Schematic illustration of FT upregulation pathways leading to AD and reversal by FT deletion or inhibition. Aberrant upregulation of FT/protein farnesylation increases the membrane association of farnesylated proteins, including Ras and Rheb, and their interactions with downstream effectors, leading to hyperactivation of mTORC1 signaling and AD. Deletion or inhibition of FT results in the reduced membrane association of Ras and Rheb and reverses the pathogenic activation of the signaling pathway.

## Author contributions

AJ performed experiments with postmortem human and mouse brain samples and cell cultures, analyzed the data, interpreted the results, and wrote the manuscript. SC performed mouse behavioral tests, ELISA, and immunoblot analysis of mouse brains. RZ quantified microglial activation in mouse brain sections and reviewed the manuscript. DAB provided postmortem human brain samples and data, discussion of the data, and critical review of the manuscript. MOB provided the floxed FT mice and suggestions on the manuscript. LL conceived the study, supervised the progress of all experiments, interpreted the results, and edited and finalized the manuscript. All authors reviewed the manuscript.

## Acknowledgments

We thank Debra Magnuson, Karen Skish, and Gregory Klein at Rush University Alzheimer’s Research Center for selecting and providing the postmortem human brain specimens, and Gunter Eckert at the University of Giessen for providing the APP695-overexpressing and control SH-SY5Y cells. We also thank Andrea Gram for breeding, maintaining and genotyping the experimental mice, Kyle LeBlanc for preparing mouse brain sections and immunostaining, and Juan E. Abrahante Lloréns at the University of Minnesota Informatics Institute for assisting with RNA-seq data analysis. AJ was partly supported by the Kwanjeong Educational Foundation Overseas Scholarship from South Korea. This work was supported in part by grants from the National Institute on Aging of the National Institutes of Health (R01AG031846, RF1AG056976, RF1AG058081, P30AG10161, and R01AG15819) and the College of Pharmacy at the University of Minnesota. ROS resources can be requested at https://www.radc.rush.edu.

## Notes

### Competing Interest Statement

The authors have declared no competing interest.

